# Identifying novel genetic and phenotypic associations to genomic features by leveraging off-target reads in exome sequencing data

**DOI:** 10.1101/2024.11.30.625754

**Authors:** Defne Ercelen, Christa Caggiano, Richard Border, Sriram Sankararaman, Serghei Mangul, Noah Zaitlen, Michael Thompson

**Affiliations:** Computational and Systems Biology Interdepartmental Program, University of California, Los Angeles, Los Angeles, CA, USA; Center for Human Genetics and Genomics, NYU Langone, New York, NY, USA; Interdepartmental Program in Bioinformatics, University of California, Los Angeles, Los Angeles, CA, USA; Department of Neurology, University of California, Los Angeles, Los Angeles, CA, USA; Department of Computer Science, University of California, Los Angeles, Los Angeles, CA 90095, USA; Department of Computational Medicine, University of California, Los Angeles, Los Angeles, CA, USA; Department of Human Genetics, University of California, Los Angeles, Los Angeles, CA, USA; Department of Computational Medicine, David Geffen School of Medicine, University of California, Los Angeles, Los Angeles, CA 90095, USA; Systems and Synthetic Biology, Centre for Genomic Regulation, Barcelona, Spain

## Abstract

Upwards of 40% of reads in sequencing datasets may be unmapped and discarded by standard protocols. Recent work has shown the utility of re-analyzing these unmapped reads to construct meaningful features, such as immune diversity repertoires or copy number variation in mtDNA and rDNA. While previous analyses of these features have produced significant correlations with diverse traits, they have generally been limited to analyses of RNA-sequencing data in phenotype-specific cohorts. Here, we explore whether associations can be identified using population-scale, whole-exome sequencing data in the UK BioBank. Using recently developed tools, we constructed multiple features including T-cell receptor diversity metrics, microbial load, and mtDNA and rDNA copy numbers for nearly 50,000 individuals in the UK BioBank. We first verify the validity of our method by showing that GWAS on these constructed traits results in replication of associations from studies in which the phenotypes were explicitly measured. Next, across several GWAS, we identified 21 novel independent significant loci in 11 genes, most of them in genes implicated in the innate immune response. Finally, we further analyzed the read-constructed features by establishing correlations to other population-level biobank traits such as immune disorders, metabolic disorders, neuropsychiatric disorders, and blood cell counts. Our results suggest that existing tools for feature construction from unmapped reads can offer novel information at the population level, and that these features can be used to establish novel genetic associations.

## INTRODUCTION

Advances in sequencing technology along with expansions in biobank availability have spurred genetic analyses of great scale and depth. Of these advances, whole-exome sequencing (WES) has emerged as a powerful tool to elucidate relationships between genetic variation and complex traits (1) (2). While high-throughput sequencing methods have enabled a wealth of discoveries, the majority of previous works that use these technologies follow protocols that disregard reads unmapped to a target reference panel. Notably, an estimated 40-60% of all exome sequencing consists of these off-target reads (3), presenting a substantial loss of potential information.

To take advantage of the entirety of sequencing reads in a dataset, researchers have developed several algorithms to remap these reads to relatively unexplored regions of the genome (4) (5). Recent work from our group and others has shown that these off-target reads can be leveraged for several purposes such as imputation of common variants and estimation of copy number variation (6) (4) (5). ImReP—one such tool that leverages reads that do not align onto the target regions in the genome—allows bulk-sequencing and off-target reads to be recycled into T-cell receptor (TCR) and B-cell receptor (BCR) profiles (7). More recently, ImReP was expanded into the Seeing Beyond the Target (SBT) platform, which, in addition to immune profiling, produces features for microbial DNA sequences, and structural variation for mitochondrial (mtDNA) and ribosomal DNA (rDNA) (8). Displaying the utility of the these off-target recycling tools, the authors were able to recapitulate cancer-related phenotype associations by profiling off-target reads from tumor-focused sequencing studies (9) (10). Nonetheless, the utility of leveraging unmapped reads from whole-exome sequencing for population-level data across a more general range of complex phenotypes remains unexplored.

Here, we address this gap by utilizing the extensive phenotypic and genetic data in the UK BioBank (UKBB) to assess the utility of off-target read recycling for large-scale studies. Using existing tools like SBT and ImReP, we derived features including mitochondrial DNA copy number, ribosomal DNA copy number, immune repertoire metrics, and microbial load from WES data for approximately 50,000 UKBB participants. Our objectives are two-fold: first, to validate the potential of recycling off-target reads as a robust method for studying underexplored genomic features by replicating known associations; and second, to demonstrate the broader utility of these features by uncovering novel genetic and phenotypic associations.

The results we present here not only reinforce the value of off-target read analysis in replicating known genetic associations but also highlight the potential to generate novel insights into diverse phenotypes such as immune disorders, metabolic traits, and neuropsychiatric conditions. By uncovering 21 new loci across 11 genes, many involved in innate immunity, we establish off-target read recycling as a useful approach for advancing population-scale genomic studies. This work underscores the importance of leveraging underutilized sequencing data to expand our understanding of the genetic architecture of complex traits, paving the way for novel discoveries and future functional studies.

## METHODS

### Data Processing and Construction Genomic Features

We downloaded the 50,000 whole-exome release in March 2019 from the UK BioBank (UKBB) consortium. Using samtools v.1.15 (11) and chromosome build hg38, we converted files from CRAM to FASTQ format. To build features of viral load, fungal load, mtDNA copy number, and rDNA (5S, 18s, and 28S) copy number, we directly input the files into Seeing Beyond the Target (SBT) (8). We used SBT with the suggested default parameters for all features (12).

From SBT, we extracted estimated copy numbers for mtDNA and rDNA (5S, 18S, and 28S), as well as counts of viral and fungal transcripts found in each sample. We refer to the features corresponding to these traits as mtDNA, rDNA5S, rDNA18S, rDNA28, viral, and fungal respectively. To avoid population structure effects while maximizing sample size, we removed non-White British and related individuals (KINGs (13) kinship coefficient greater than 0.0442, 3rd degree). Further, we removed outliers from each constructed feature (defined by having a phenotypic value beyond 2 standard deviations of the mean), which resulted in n=45,242 (mtDNA), n=44,923 (rDNA5S), n=43,784 (rDNA18S), n=44,791 (rDNA28S), n=44,556 (viral), n=44,640 (fungal) samples for each feature. Histograms of the distribution of each feature after filtering can be seen in **Figure 1** for TCRA features and **Supplementary Figure 1** for SBT features.

**Figure 1:**
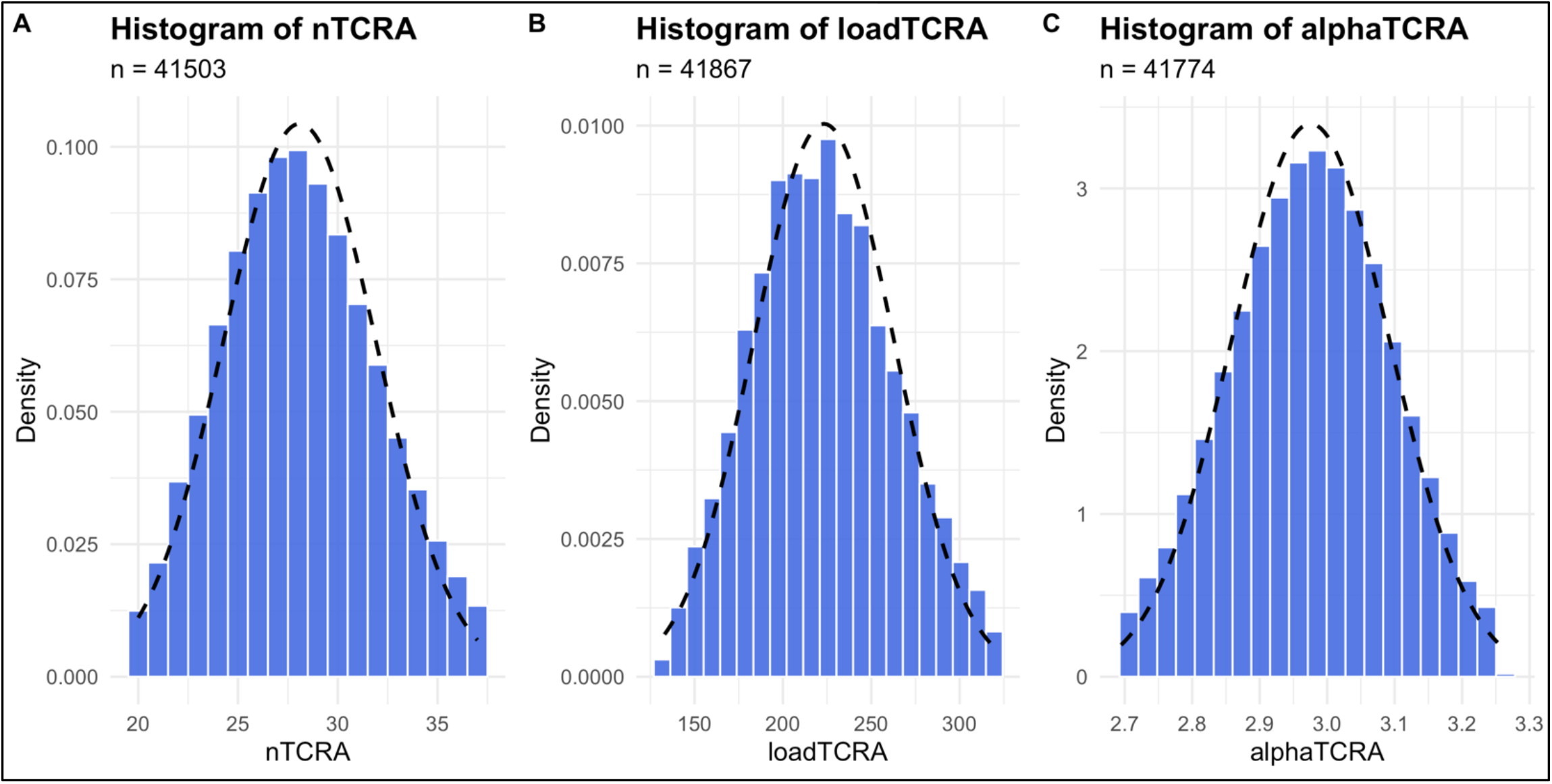
Histograms of TCRA feature distributions after QC (8). Histograms for other features can be found in Supplementary Figure 1. (A) Distribution of feature nTCRA, measured as each distinct TCRA sequence found per sample. (B) Distribution of feature loadTCRA, measured as each partial TCRA sequence found per sample. (C) Distribution of feature alphaTCRA, measured as the alpha diversity (Shannon entropy) of TCRA sequences per sample.

To build the immune repertoire features using ImReP, we followed a protocol published in previous reports (14). Namely, we trimmed read files to regions of immune-related transcripts and to regions with low coverage (less than 10x) to maximize potential targets while remaining computationally efficient (14). We passed low coverage sequences into ImReP to account for immune-sequence diversity. Many alignment algorithms use majority voting to reach a definitive alignment for a sequence (15). Since immune-related sequences, such as VDJ regions, show high genetic diversity between samples, they usually have low agreement in the alignment algorithm, and thus low coverage (16). This attribute makes these regions viable targets for off-target reads for tools such as ImReP, which leverage collections of unmapped reads (14).

From ImReP, we obtained the amino acid sequences of each B-cell receptor (BCR) and T-cell receptor (TCR) transcript, the chain type (antibody chain type for BCR: H, K, L; and for TCR: alpha, beta, delta, gamma), the aggregate number of reads fully or partially mapped to each transcript, and the number of different of VDJ sequences in each sample. We then further processed this data into diversity measures of BCR and TCR transcripts for each sample (17). Notably, over 90% of samples had counts of 0 for all immune chains except for TCR alpha (TCRA). This issue likely arises due to limitations in the capture kit used for sequencing, as it lacks a sufficient number of probes specifically designed for amplifying BCR/TCR regions; for example, omitting crucial T cell genes like TRBD1, TRBD2, TRDC, and TRGC2 (18). We therefore chose to only use TCRA features in downstream analyses. Henceforth, we refer to the phenotype for the number of each distinct TCRA sequence as “nTCRA”, the phenotype for the number of partial TCRA sequences as “loadTCRA”, and the phenotype for the alpha diversity of TCRA sequences per sample as “alphaTCRA”. We performed the same quality control and filtering steps as mentioned above, resulting in n=41,503 (nTCRA), n=41,868 (loadTCRA), n=41,774 (alphaTCRA) samples for each phenotype.

### Genome-Wide Association Studies

We used Plink2 (19) to run all genome-wide association studies (GWAS). In addition to excluding non-White British, related individuals, and outliers from the analyses as described above, we filtered for single nucleotide polymorphisms (SNPs) with a minor allele frequency of 0.001, INFO score 9, and Hardy-Weinberg equilibrium at threshold p=1E-10. Explicitly, we ran all GWAS with parameters –geno 0.05, –hwe 1e-10, –maf 0.001, and with sex, age, sexXage, age^2^, as well as the first 20 genotypic principal components (PCs) computed on the filtered dataset as covariates on 7,682,621 SNPs. When running GWAS on the TCRA phenotypes, we also included white blood cell counts as covariates to control for cell composition.

### Complex phenotype associations with features

To relate the constructed phenotypes with other complex traits in the UKBB, we conducted association tests using linear and logistic models, corresponding to continuous and binary phenotypes respectively. For each phenotype, we ran a regression with the complex trait as the outcome variable and an SBT-constructed feature, sex, age, sexXage, age^2^, the first 20 principal components (PCs), as well as white blood cell count as covariates. We ran the association tests for each respective feature from the GWAS study, leading to roughly 41K individuals (exact numbers above) for each association test. We controlled for multiple testing by using Bonferroni adjustment.

To validate the SBT-constructed features, we sought to replicate associations between mtDNA copy number and kidney-related phenotypes such as platelet crit, creatinine in urine, cystatin c, and urea as reported in a previous paper (20) (Table 1). After validating the mtDNA SBT feature, we sought to discover novel associations of our features with complex phenotypes available in the UKBB including immune, metabolic, and mental disorders, and white blood counts (in this test we exclude white blood cell counts as covariates). The following complex phenotypes were tested for all constructed features: asthma, hay fever/allergy, depression, overall health, depression of a parent, Alzheimer’s disease/dementia of a parent, and white blood cell counts (leukocyte, lymphocyte, monocyte, neutrophil, eosinophil, basophil). All tested binary complex phenotypes had a prevalence of at least 5% in the sample population.

**Table 1:** Phenotypes trom Yonova-Doing et al. (20) that were tested against mtDNA copy number. All association tests were significant after Bonferroni correction. Effect sizes are in the same direction as the previous study.

| ***Phenotype*** | **mtDNA CN** |  |  |
| --- | --- | --- | --- |
|  | *Effect Size* | *Std. Error* | *P value* |
| *Platelet crit* | 2.652E-3 | 6.06E-4 | **1.19E-5** |
| *Creatinine in Urine* | -3.368E-2 | 7.12E-1 | **2.27E-6** |
| *Cystatin c* | -2.015E-2 | 2.19E-3 | **< 2E-16** |
| *Urea* | -0.0987420 | 1.78E-2 | **3.19E-8** |

## RESULTS

### Overview of constructed features

After running the SBT platform with recommended parameters and doing relevant quality control (QC) and filtering, we arrived at sample sizes of roughly 44.5K for each SBT feature (See Methods for exact numbers). The ranges, means and standard deviations (SD) are as follows: mtDNA 0-1.93, 0.88 (0.38); viral 17-695, 349.5 (115.10); fungi 0-6, 2.76 (1.64); rDNA5S 0-5.12, 2.28 (1.03); rDNA18S 0-4.96, 2.37 (0.95); rDNA28S 0-9.03, 4.25 (1.74). We then applied the same QC and filtering to our ImReP pipeline to construct features representing immunologic TCRA profiles. Roughly 41K individuals passed all steps, with ranges and mean values (SD) for each feature as follows: nTCRA 20-37, 28.09 (3.82, loadTCRA 133-321, 223.2 (39.77), alphaTCRA 2.695-3.253(0.12; **Figure 1**).

### Replication of features through previous studies of mtDNA

To first establish the validity of the constructed features, we performed a replication analysis. Here, we sought to replicate the associations of Yonova-Doing et al., which established associations between kidney-related phenotypes and mtDNA copy number (20). We replicated these previous associations with our method with high statistical power (*p*<0.001, and effect size in the same direction for each test; **Table 1**).

### Replication and discovery in genome-wide association studies

We next conducted genome-wide association studies (GWAS) for each of our features using imputed genotyping data from the UKBB on 7,682,621 SNPs. The sample sizes for each of the features were as follows: nTCRA 41,503; loadTCRA 41,867; alphaTCRA 41,774; mtDNA 45,242; rDNA5S 44,923; rDNA18S 43,784; rDNA28S 44,791; viral 44,556; and fungal 44,640. We identified 21 independent novel significant loci (p-value < 10e-8) in the GWAS of the TCRA, mtDNA, and rDNA features (Table 2). Within each locus, we report the SNPs with lowest p-values and highlight the gene containing these SNPs (except for the three SNPs noted “intergenic”).

**Table 2:** GWAS summary statistics for significant independent loci. We report the SNP with the lowest p-value within a +-1 Mb window and its nearest gene. Check data availability statement for full GWAS summary statistics. Genetic loci for hits were determined as 1 Mbp +/-from the SNP with the lowest p-value. ADAM11 and LINC01252 had significant hits in all TCRA GWAS, followed by PLEKHB2 and ACAP2, which had significant hits in alphaTCRA and nTCRA GWAS

| Feature | Chrom. | Position | ID | Effect Size | Std. Error | P value | Gene |
| --- | --- | --- | --- | --- | --- | --- | --- |
| alphaTCRA | 2 | 131150260 | rs146762588 | 2.26E-2 | 1.55E-3 | 2.55E-48 | PLEKHB2 |
| alphaTCRA | 3 | 195335589 | rs9886975 | 5.44E-3 | 9.51E-4 | 1.04E-8 | ACAP2 |
| alphaTCRA | 6 | 32628428 | rs2854275 | -8.17E-3 | 1.33E-3 | 8.44E-10 | HLA-DQA1 |
| alphaTCRA | 6 | 31604591 | rs10885 | -9.34E-3 | 1.19E-3 | 3.41E-15 | intergenic |
| alphaTCRA | 12 | 11556951 | rs150588766 | 2.78E-2 | 2.91E-3 | 1.05E-21 | LINC01252 |
| alphaTCRA | 17 | 44781030 | rs199440 | 1.32E-2 | 1.23E-3 | 4.95E-27 | ADAM11 |
| alphaTCRA | 17 | 43673792 | rs62064441 | 1.23E-2 | 1.16E-3 | 1.76E-26 | LINC02594 |
| nTCRA | 2 | 131158755 | rs145650663 | 5.67E-1 | 5.14E-2 | 2.81E-28 | PLEKHB2 |
| nTCRA | 3 | 195335589 | rs9886975 | 1.75E-1 | 3.16E-2 | 3.31E-8 | ACAP2 |
| nTCRA | 6 | 31597700 | rs3130623 | -2.23E-1 | 3.92E-2 | 1.22E-8 | intergenic |
| nTCRA | 12 | 11556951 | rs150588766 | 6.22E-1 | 9.68E-2 | 1.35E-10 | LINC01252 |
| nTCRA | 17 | 43656176 | rs1724438 | 4.24E-1 | 4.03E-2 | 6.45E-26 | MEOX1 |
| nTCRA | 17 | 44781030 | rs199440 | 4.22E-1 | 4.09E-2 | 6.03E-25 | ADAM11 |
| loadTCRA | 2 | 131504831 | rs79058854 | 4.14E+00 | 5.24E-1 | 2.89E-15 | pseudo gene<br>SMPD4BP |
| loadTCRA | 12 | 11557536 | rs73287481 | 6.49E+00 | 9.93E-1 | 6.26E-11 | LINC01252 |
| loadTCRA | 12 | 52694533 | rs11170081 | -2.42E+00 | 4.41E-1 | 4.31E-8 | KRT77,<br>ENSG00000257700 |
| loadTCRA | 17 | 43573061 | rs62065452 | -5.36E+00 | 3.82E-1 | 1.08E-44 | DHX8 |
| loadTCRA | 17 | 44773783 | rs199441 | -4.81E+00 | 3.87E-1 | 2.03E-35 | ADAM11 |
| rDNA5S | 1 | 228601638 | rs472594 | 3.65E-1 | 8.82E-3 | 1.49E-367 | intergenic |
| rDNA5S | 1 | 227249587 | rs72748155 | 3.47E-1 | 1.92E-2 | 1.13E-72 | CDC42BPA |
| rDNA5S | 1 | 226193200 | rs143974947 | 1.92E-1 | 1.96E-2 | 1.55E-22 | LOC101927247 |

We examined the validity of our GWAS results using a previous GWAS of measured mtDNA copy number that utilized approximately 200,000 genomes from the UK BioBank (21). Although the sample size in the present study comprised only 25% of the previous study (∼50,000) and employed whole-exome sequencing rather than whole-genome sequencing, we achieved significance (p-value < 0.05) at 32.8% of their significant SNPs within our GWAS. This demonstrates that even with smaller cohorts and less comprehensive sequencing, it is possible to identify highly significant associations. Moreover, these results paired with the replication of the associations with Yonova-Doing et al. suggest that our constructed features indeed capture signal of their intended phenotype.

After validating our GWAS results, we next examined several of the implicated loci. First, we note that the *HLA-DQA1* (effect size -8.17×10^−8^, p-value 8.44×10^−10^) locus was significant in the alphaTCRA GWAS, again, suggesting that our approach does indeed capture the intended TCR alpha chain genes. Further, loci in genes *PLEKHB2* (2.26×10^−2^, 2.55×10^−48^) and *ACAP2* (5.44×10^−^ 3, 1.04×10^−8^) were significant in the alphaTCRA and nTCRA GWAS, and a locus within the gene *ADAM11* (1.32×10^−2^, 4.95×10^−27^) was significant in all three TCRA GWAS. While *ADAM11* was significant across all three TCRA GWAS, we were unable to find previous literature suggesting a role in immune-related pathways. The current suggested role for *ADAM11* is as a non-proteolytic metalloprotease which plays a role in synaptic transmission and neurogenesis (22). We did not discover any loci at genome-wide significance in the rDNA18S, rDNA28S, mtDNA, viral or fungal GWAS. Manhattan plots for each GWAS can be seen in **Figure 2** and **Supplementary Figure 2** and QQ plots for all features can be seen in **Supplementary Figure 3**.

**Figure 2:**
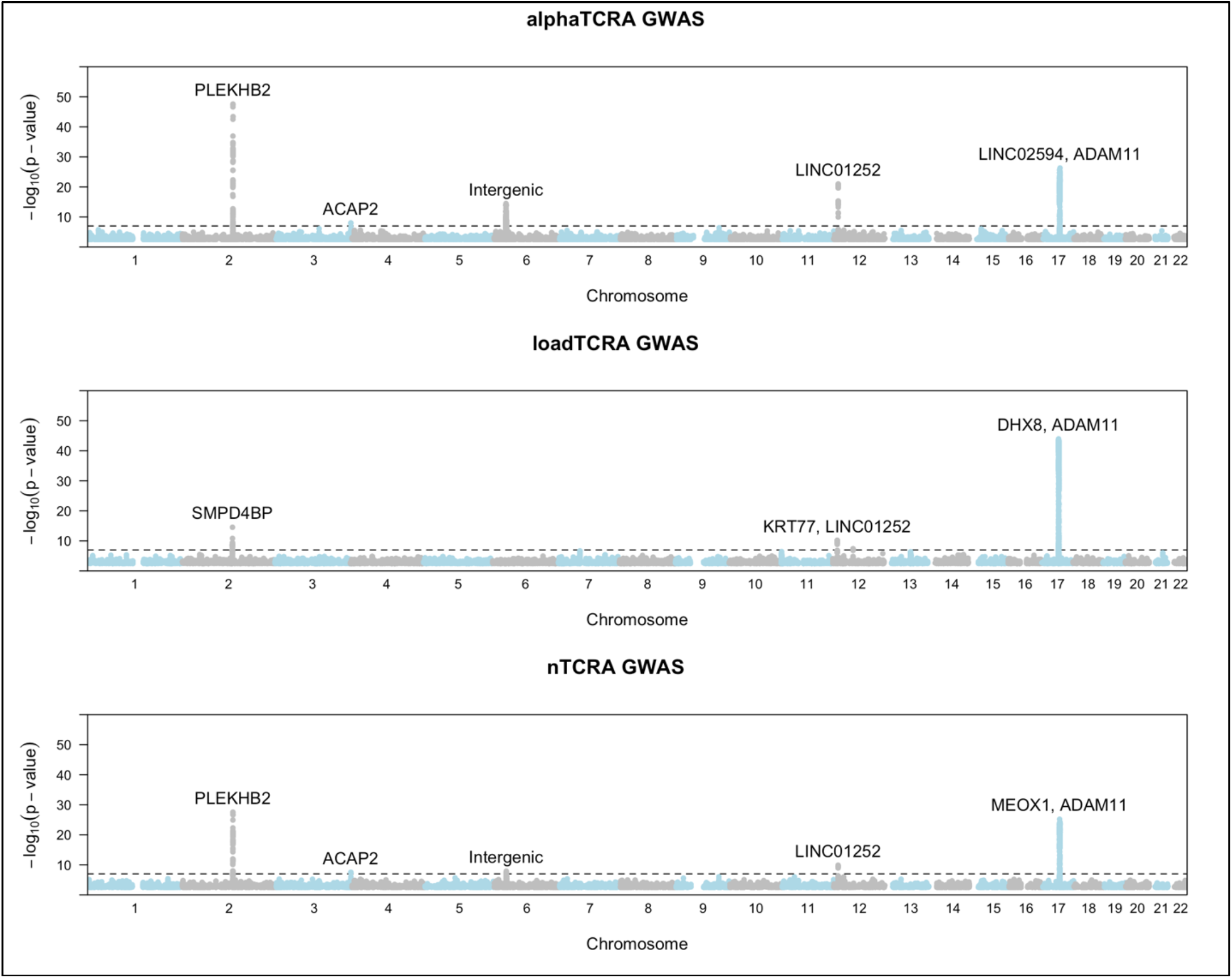
Manhattan plots of TCRA GWAS alphaTCRA, loadTCRA, nTCRA. Genes with hits at the highest -log(p-values) are labeled. The dashed line indicates a -log(p-value) threshold of 1×10-8. Top hits are observed in chromosomes 2,12, and 17, with other, weaker hits on chromosomes 3 and 6. Samples sizes are as follows: alphaTCRA (n=41,774), loadTCRA (n=41,867), nTCRA (n=41,503). The rest of the GWAS can be found in Supplementary Figure 2.

Many of the genes that were significant in all TCRA GWAS were implicated in T-cell cycle regulation and antigen presentation (23) (24). Out of these genes *MEOX1* (p=1.11118×10^−39^) and *PLEKHB2* (p=6.22×10^−44^) contained many of the strongest hits in the alphaTCRA and nTCRA GWAS, respectively. Other genes of interest were *HLA-DQA1, ACAP2, DHX8*, which play significant roles in inflammatory responses, natural cytotoxicity, and RNA splicing, respectively (25) (26) (27).

Numerous significant associations were identified on chromosome 1 in the rDNA5S GWAS, which aligns with biological expectations, as all genes encoding the 5S ribosomal subunit are located on this chromosome (28) (29). R Successfully recapitulating this known biology not only underscores the robustness of our findings but also serves as a validation of our methodology for accurately capturing and characterizing rDNA features.

### Feature associations with other complex phenotypes

We next investigated the relationship between our constructed features and several related disorders and environmental factors. We ran association tests with the SBT-constructed features and multiple complex phenotypes including immune disorders, metabolic disorders, neuropsychiatric disorders, blood counts, and illnesses of parents. We ran linear and logistic regressions depending on the phenotype (linear for continuous, logistic for binary) with the outcome as the complex phenotype (Methods). The full list of complex phenotypes tested and summary of association tests can be seen in **Table 3** for significant associations and **Supplementary Table 1** for the non-significant features.

**Table 3:** Phenotype associations across features that yielded significant associations. For the non-significant associations refer to Supplementary Table 2. Continuous traits were tested using linear regression and binary traits were tested using logistic regression. Bolded p-values represent significance after Bonferroni correction.

| Phenotype | rDNA(5S) |  |  | rDNA(18S) |  |  | rDNA(28S) |  |  | mtDNA |  |  |
| --- | --- | --- | --- | --- | --- | --- | --- | --- | --- | --- | --- | --- |
|  | Effect Size | Std. Error | P value | Effect Size | Std. Error | P value | Effect Size | Std. Error | P value | Effect Size | Std. Error | P value |
| Age | -4.90E-02 | 3.89E-02 | 2.08E-01 | -2.46E-02 | 4.29E-02 | 5.66E-01 | -4.06E-02 | 2.30E-02 | 7.76E-02 | -7.38E-01 | 1.04E-01 | **1.59E-12** |
| Asthma | -6.98E-04 | 2.02E-03 | 7.30E-01 | 3.14E-03 | 2.22E-03 | 1.58E-01 | 2.99E-03 | 1.54E-03 | 4.31E-02 | -4.36E-03 | 5.45E-03 | 4.24E-01 |
| Hay fever/Allergic Rhinitis | -1.82E-03 | 1.52E-03 | 2.32E-01 | -3.45E-03 | 1.68E-03 | 3.99E-02 | -1.01E-03 | 8.96E-04 | 2.59E-01 | 6.66E-04 | 4.08E-03 | 8.70E-01 |
| Depression | 4.99E-04 | 1.49E-03 | 7.37E-01 | 3.50E-03 | 1.64E-03 | 3.32E-02 | 4.80E-04 | 8.75E-04 | 5.84E-01 | 2.81E-03 | 4.00E-03 | 4.81E-01 |
| Basophil Count | 8.60E-04 | 2.56E-04 | **7.75E-04** | 1.07E-03 | 2.78E-04 | **1.21E-04** | 6.31E-04 | 1.49E-04 | **2.17E-05** | -2.65E-03 | 6.84E-04 | **1.09E-04** |
| Eosinophil Count | -6.90E-04 | 6.49E-04 | 2.88E-01 | -2.09E-04 | 7.14E-04 | 7.70E-01 | -2.65E-04 | 3.83E-04 | 4.89E-01 | -9.23E-03 | 1.74E-03 | **1.19E-07** |
| Lymphocyte Count | 6.02E-03 | 5.54E-03 | 2.77E-01 | -4.33E-04 | 6.13E-03 | 9.44E-01 | 2.33E-03 | 3.34E-03 | 4.85E-01 | 2.69E-02 | 1.43E-02 | 6.05E-02 |
| Monocyte Count | 1.80E-03 | 9.89E-04 | 6.88E-02 | 2.98E-03 | 1.09E-03 | 6.11E-03 | 1.81E-03 | 5.82E-04 | **1.82E-03** | -1.81E-02 | 2.66E-03 | **9.80E-12** |
| Neutrophil Count | -2.42E-03 | 6.82E-03 | 7.22E-01 | 1.38E-02 | 7.51E-03 | 6.69E-02 | 5.94E-03 | 4.03E-03 | 1.40E-01 | -6.19E-01 | 1.80E-02 | **< 2E-16** |
| White Blood Cell Count | 9.26E-03 | 1.02E-02 | 3.65E-01 | 1.72E-02 | 1.13E-02 | 1.28E-01 | 1.08E-02 | 6.07E-03 | 7.48E-02 | -6.21E-01 | 2.64E-02 | **< 2E-16** |
| Overall health Rating | 2.98E-03 | 3.62E-03 | 4.10E-01 | 1.20E-02 | 3.98E-03 | **2.65E-03** | 9.52E-03 | 2.13E-03 | **8.15E-06** | -4.45E-02 | 9.70E-03 | **4.54E-06** |
| Alzheimer's/ Dementia of mother | 3.30E-03 | 1.34E-03 | 1.39E-02 | 4.41E-03 | 1.48E-03 | **2.85E-03** | 3.40E-03 | 7.94E-04 | **1.89E-05** | 7.32E-03 | 3.63E-03 | 4.39E-02 |
| Severe Depression of mother | 9.90E-04 | 1.19E-03 | 4.05E-01 | 1.25E-03 | 1.31E-03 | 3.41E-01 | 1.13E-03 | 7.02E-04 | 1.08E-01 | -2.37E-03 | 3.20E-03 | 4.58E-01 |

In addition to our initial replication analysis of kidney-related phenotypes and mtDNA, we found further relationships of mtDNA with other phenotypes. mtDNA copy number was significantly associated (at Bonferroni-adjusted threshold < 0.003) with almost all blood counts min. p-value < 2×10^−^16), as well as age (p-value 1.59×10^−12^) and overall health (p-value 4.54×10^−6^) (Table 3). We hypothesize this association may be due to recent evidence that mitochondrial DNA variation and integrity are linked to aging and age-related health outcomes (30) (31) (32) (33) (34) (35).

Additional phenotype associations that passed our p-value threshold (Bonferroni correction, p-value < 0.003) and were significantly associated to rDNA features were Alzheimer’s disease/dementia of mother (min. p-value 1.89×10^−5^), basophil count (min. p-value 2.17×10^−5^), and overall health rating (min. p-value 8.15×10^−6^, Table 3). We did not find any significant associations of TCRA features and viral or fungal load.

## DISCUSSION

In this manuscript, we explore at population scale the utility of tools leveraging off-target reads for the construction of features related to understudied and hypervariable genomic regions, such as immune-related genes, copy number variation in ribosomal and mitochondrial DNA, and microbial genome load. Using data from the UK BioBank, we show that features derived from these tools recapitulate known biology, can be used in GWAS, and can be examined through additional tests of association for discovery of novel relationships.

We validate the phenotypes constructed from the off-target reads via replication of two previous, independent studies. We first replicate associations of mtDNA copy number (mtDNACN) with several kidney and excretory-system-related phenotypes, including platelet crit (p-value 1.19×10^−^ 5), creatinine in urine (p-value 2.27×10^−6^), cystatin c (p-value <2×10^−16^), and urea (p-value 3.19×10^−^ 8) levels (20). Moreover, in line with the relation of mtDNACN variability in hematological disorders, we found significant associations between mtDNACN and blood counts of basophils (p-value 1.09×10^−4^), eosinophils (p-value 1.19×10^−7^), monocytes (p-value 9.80×10^−12^), neutrophils (p-value <2×10^−16^), and overall white blood cell count (p-value <2×10^−16^) (21) (36). Separately, we replicated a collection of variants discovered in a GWAS of measured mtDNA copy number that used a substantially larger sample size, providing further evidence for the validity of our approach.

Our GWAS over the constructed features revealed 21 significant loci associated with immune diversity at genome-wide significance. Notably, several loci in key immune genes, including *PLEKHB2* (min. p-value 2.55×10^−48^), *ADAM11* (min. p-value 2.03×10^−35^), *ACAP2* (min. p-value 1.04×10^−8^), *HLA-DQA1* (p-value 8.44×10^−10^), *MEOX1* (p-value 6.45×10^−26^), *DHX8* (p-value 1.08×10^−44^), and *KRT77* (p-value 4.31×10^−8^), were identified. *MEOX1, PLEKHB2*, and *ADAM11* emerged as recurrent hits across all TCRA features, suggesting their potential as novel markers for immune diversity. *MEOX1* functions as a transcription factor crucial for T-cell migration and proliferation, while *PLEKHB2* is implicated in retrovirus binding, along with prior links to influenza vaccine efficacy (37) (38) (39) (40) (41) (23) (24) (42). Additionally, *ACAP2* is associated with vaccine response modulation, particularly in host-pathogen interactions with Vaccinia virus (43) (44) (45). These findings may potentially unveil novel candidate genes for immune compartment and antigen receptor diversity. We note that while our discoveries may spur biological interest, they will require further scrutiny, as the phenotypes we used may be noisy realizations of the phenotypes they are intended to capture.

Considering that regulation of structural variation in ribosomal DNA has not been well-characterized, we examined copy number variation in rDNA for different ribosomal subunits 5S, 18S, and 28S. In line with previous reports that structural variation in rDNA is known to vary between humans by 2 orders of magnitude and to exhibit tissue and disease specificity, we show a wide range of rDNA copy number variation in all three of our rDNA features (46) (47). Our approach suggests the potential of using existing exome datasets to characterize rDNA and its regulation in further studies. We also found that structural variation in mitochondrial and ribosomal DNA was associated with overall health (min p-value 8.15×10^−6^). This result is consistent with observations that mtDNA copy number is known to be important in maintaining healthy physiology (21). In addition many there have been many recent works implicating mitochondrial variants’ effects in diseases spanning many organ systems such as cardiac, metabolic, and excretory (20) (21) (48) (49) (50) (51). However, rDNA copy number is a relatively understudied feature, and there is unexplored potential in studying the associations of rDNA variation and population or ecological density (47).

We observed no significant associations of microbial genome load on any of our GWAS or complex trait associations. One reason for this might be that whole exome sequencing (WES) of genomic DNA is not a good source for this type of data. This is a foreseen limitation of our study as the industry gold standard for microbial sequencing is 16s RNA or metagenomic sequencing, which utilizes a whole different protocol than genomic WES (52).

We are hopeful that our analyses here will encourage the use of off-target tools for large-scale whole-exome sequencing datasets in either direct, relevant traits of interest, or as covariates in downstream association analyses. Finally, as biobanks increasingly adopt whole-exome sequencing for their cohorts, the work presented here may be expanded across diverse populations, including All of Us, FinnGen, Biobank Japan, and a new African biobank 54Gene (53) (54) (55) (56), spurring novel discovery.

## DATA AVAILABILITY

Individual-level sequence and phenotype data utilized in this study were obtained from the UK Biobank under application number 33127. Instructions for access to UK Biobank data are available at https://www.ukbiobank.ac.uk/enable-your-research.

Summary statistics for the GWAS conducted in this study have been deposited to Zenodo and are freely available at https://doi.org/10.5281/zenodo.14210828.

## Notes

https://doi.org/10.5281/zenodo.14210828.

### Competing Interest Statement

The authors have declared no competing interest.

