## Supplementary Figures for "Identifying novel genetic and phenotypic associations to genomic features by leveraging off-target reads in exome sequencing data"

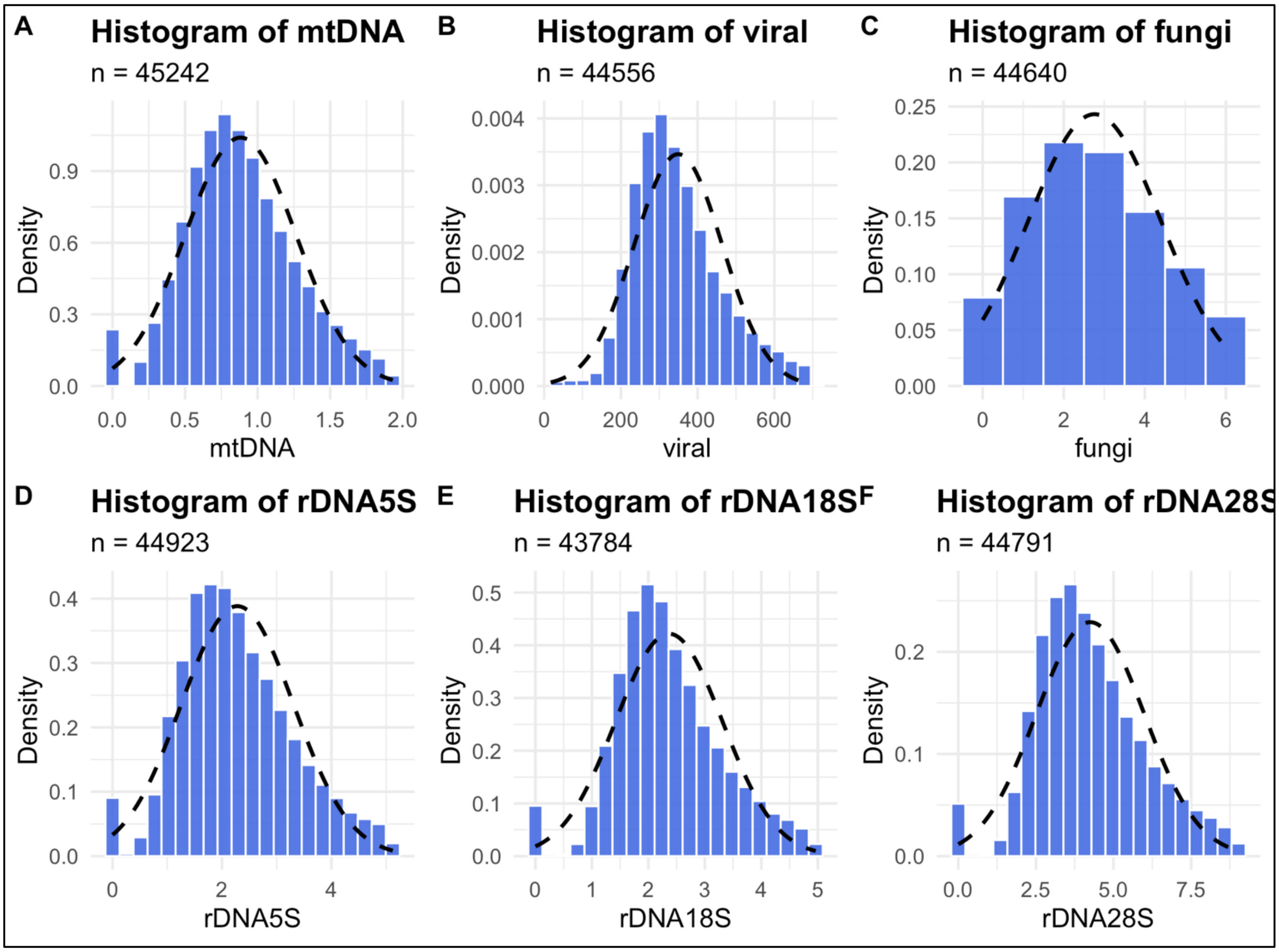


**Supplementary Figure 1:** Histograms of genomic and microbial feature distributions after QC. All measurements are adjusted for off-target coverage. (a) Distribution of feature mtDNA, measured based on reads covering mtDNA genome. (b) Distribution of feature viral, measured based on the number of reads mapped to viral genomes. (c) Distribution of feature fungi, measured based on the number of reads mapped to fungal genomes. (d) Distribution of feature rDNA5S, measured based on the reads covering 5S rDNA region. (e) Distribution of feature rDNA18S, measured based on the reads covering 18S rDNA region. (f) Distribution of feature rDNA28S, measured based on the reads covering 28S rDNA region.

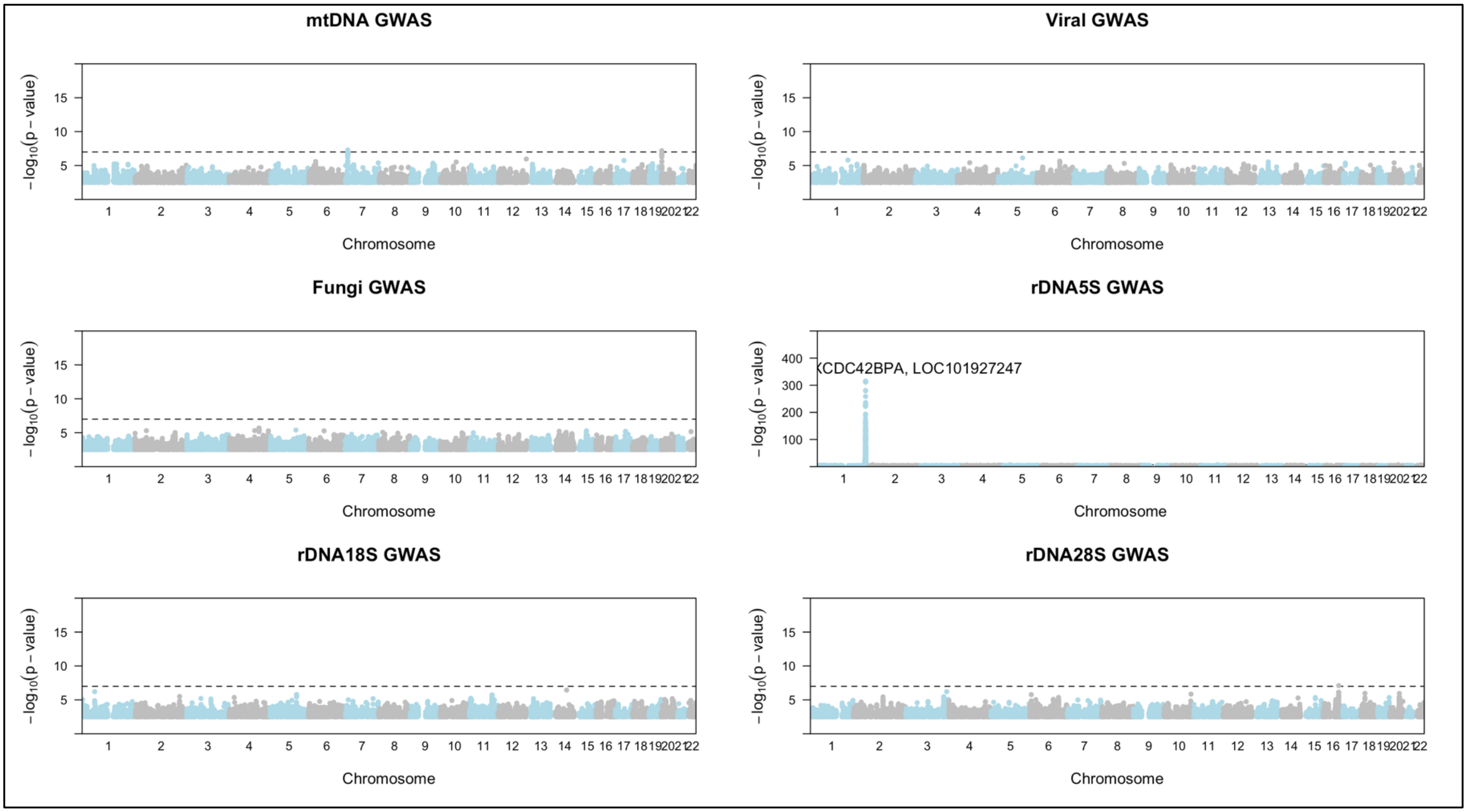


**Supplementary Figure 2:** Manhattan plots of TCRA GWAS. Genes with hits at the highest -log(p-values) are labeled. Top hits are observed in chromosome 1 for the rDNA5s GWAS, with some marginally significant hits on chromosome 6. The dashed line indicates a -log(p-value) threshold of $1x10^{-8}$. Samples sizes are as follows: (a) mtDNA (n=45,242), (b) viral (n=44,556), (c) fungi (n=44,6400), (d) rDNA5S (n=44,923), (e) rDNA18s (n=43,784), (f) rDNA28S (n=44,791). TCRA feature GWAS can be found in Figure 2.

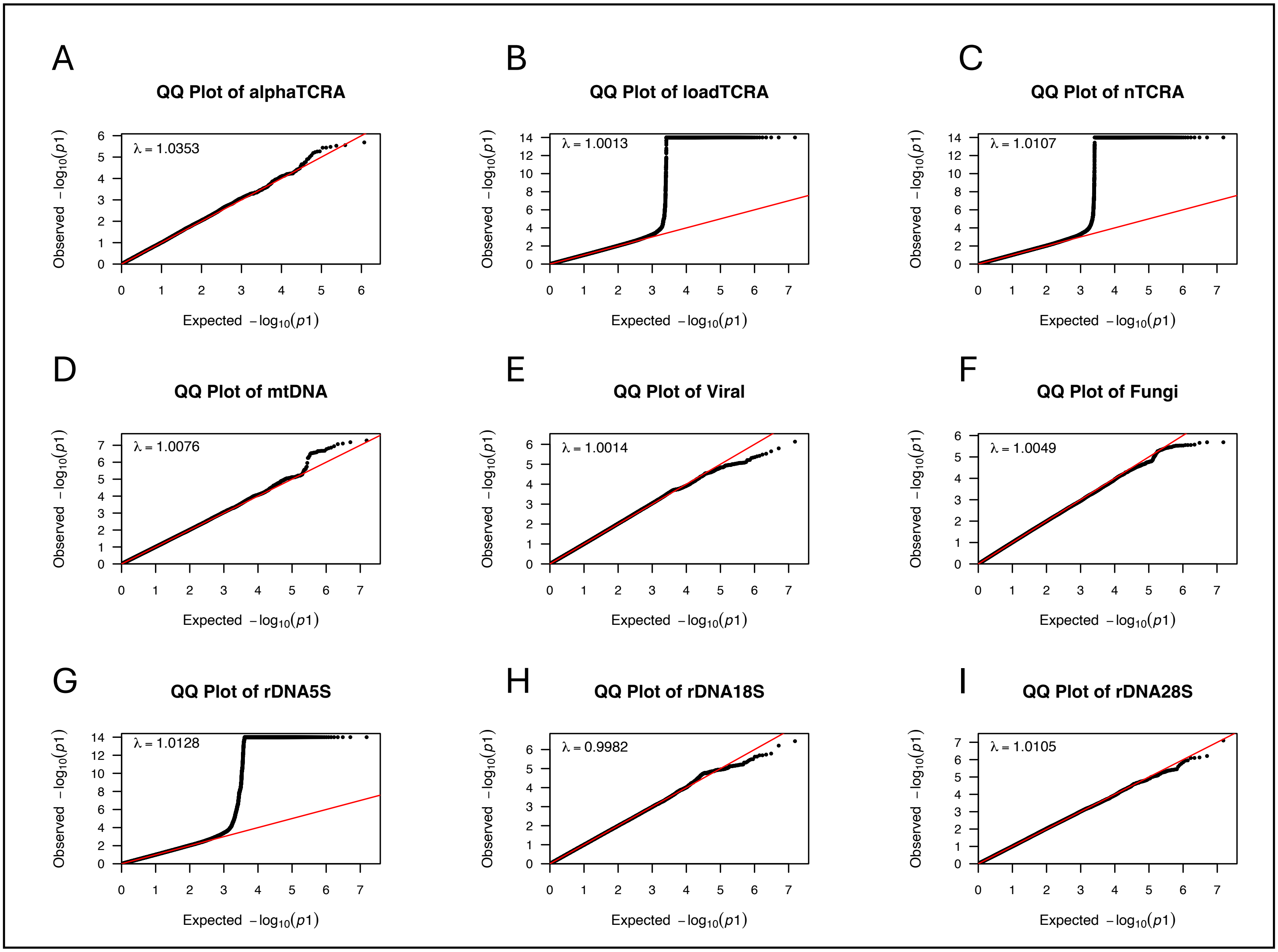


**Supplementary Figure 3**: QQ plots for all features. Significant deviations from the expected x=y line for GWAS of features loadTCRA, nTCRA and rDNA5s (panels B, C, and G). For panels B and C, these deviations are caused by the SNPs that are in heavy linkage disequilibrium with the GWAS hits. For panel G this can be explained by the fact that the entirety of rDNA 5S are localized on chromosome 1 (55) (56). For all panels the majority of the SNPs fall into the left side of the plots, meaning that they correlate with the expected -log(p-value).

| Phenotype | alphaTCRA | | | loadTCRA | | | nTCRA | | | Viral | | | Fungal | | |
|---|---|---|---|---|---|---|---|---|---|---|---|---|---|---|---|
| | *Effect Size* | *Std. Error* | *P value* | *Effect Size* | *Std. Error* | *P value* | *Effect Size* | *Std. Error* | *P value* | *Effect Size* | *Std. Error* | *P value* | *Effect Size* | *Std. Error* | *P value* |
| Asthma | -1.07 | 0.71 | 0.13 | -3.11E-03 | 2.03E-03 | 0.13 | -0.05 | 0.02 | 0.032 | *-1.35E-05* | *1.82E-05* | 0.46 | *-1.42E-03* | *1.26E-03* | 0.26 |
| Hay fever/Allergic Rhinitis | 0.94 | 0.68 | 0.16 | -1.90E-03 | 2.02E-03 | 0.35 | 0.03 | 0.02 | 0.11 | *-1.07E-05* | *1.37E-05* | 0.43 | *-2.49E-04* | *9.43E-04* | 0.80 |
| Depression | -1.05 | 0.51 | 0.04 | 2.04E-05 | 1.50E-03 | 0.99 | -0.01 | 0.02 | 0.40 | *1.01E-05* | *1.34E-05* | 0.45 | *-3.71E-04* | *9.26E-04* | 0.69 |
| Basophil Count | 4.22E-03 | 2.12E-03 | 0.05 | -1.88E-06 | 6.20E-06 | 0.76 | 1.69E-04 | 6.53E-05 | 9.54E-03 | *-2.29E-06* | *2.31E-06* | 0.32 | *-3.14E-04* | *1.59E-04* | 4.82E-02 |
| Eosinophil Count | -3.31E-03 | 5.36E-03 | 0.53 | -2.14E-05 | 1.59E-05 | 0.18 | -2.35E-04 | 1.67E-04 | 0.12 | *-8.76E-06* | *5.86E-06* | 0.14 | *-1.84E-04* | *4.04E-04* | 0.65 |
| Lymphocyte Count | -5.59E-03 | 0.03 | 0.84 | -3.63E-05 | 9.73E-05 | 0.71 | -2.87E-04 | 1.04E-03 | 0.78 | *4.69E-05* | *5.13E-05* | 0.36 | *-7.07E-03* | *3.54E-03* | 4.59E-02 |
| Monocyte Count | -0.01 | 7.73E-03 | 0.09 | -2.55E-05 | 2.28E-05 | 0.27 | -2.22E-04 | 2.40E-04 | 0.35 | *4.87E-06* | *8.96E-06* | 0.59 | *-4.17E-04* | *6.27E-04* | 0.51 |
| Neutrophil Count | 0.02 | 0.03 | 0.52 | 8.09E-05 | 1.00E-04 | 0.42 | 6.59E-04 | 1.07E-03 | 0.78 | *3.77E-06* | *6.16E-05* | 0.95 | *1.33E-03* | *4.25E-03* | 0.76 |
| White Blood Cell Count | 0.02 | 0.08 | 0.77 | 1.10E-04 | 2.51E-04 | 0.66 | 8.76E-04 | 2.67E-03 | 0.74 | *5.67E-05* | *9.32E-05* | 0.54 | *-4.47E-03* | *6.42E-03* | 0.49 |
| Overall health Rating | 0.01 | 0.03 | 0.70 | 5.60E-04 | 9.10E-05 | 4.30E-03 | 1.59E-03 | 9.55E-04 | 0.01 | *-1.78E-05* | *3.26E-05* | 0.59 | *2.73E-03* | *2.25E-03* | 0.22 |
| Alzheimer's/ Dementia of mother | -0.02 | 0.16 | 0.90 | -2.86E-04 | 4.58E-04 | -0.62 | -0.003 | 0.005 | 0.56 | *-2.05E-07* | *1.22E-05* | 0.99 | *-3.13E-04* | *8.43E-04* | 0.71 |
| Severe Depression of mother | -0.11 | 0.16 | 0.50 | -3.85E-04 | 4.60E-04 | 0.40 | -0.004 | 0.005 | 0.44 | *7.90E-06* | *1.07E-05* | 0.462 | *-1.27E-03* | *7.41E-04* | 8.65E-02 |

**Supplementary Table 1:** Phenotype associations across features that yielded non-significant associations. For the significant associations refer to **Table 3**. Continuous traits were tested using linear regression and binary traits were tested using logistic regression.
